# hSpindly’s dynamic controls SAC activity independently of the KBB pathway at unattached kinetochores

**DOI:** 10.1101/2025.07.25.666731

**Authors:** Mar Mora-Santos, Joaquín Herrero-Ruiz, Dirk Fennema-Galparsoro, Alejandro Belmonte-Fernández, Nuria Ferrandiz, Francisco Romero

## Abstract

The Spindle Assembly Checkpoint (SAC) ensures that sister chromatids do not separate until all chromosomes are properly attached to spindle microtubules and correctly bi-oriented. SAC controls the metaphase-to-anaphase transition by inhibiting the Anaphase Promoting Complex/Cyclosome (APC/C) through the formation of the Mitotic Checkpoint Complex (MCC). A critical step in this process is the recruitment of the Mad1–Mad2 complex to unattached kinetochores. Two pathways are known to mediate this recruitment: the KBB pathway (KNL1–Bub3–Bub1) and the RZZ pathway (Rod– Zw10–Zwilch). Here, we demonstrate that hSpindly plays a central role in controlling the recruitment of the Mad1–Mad2 complex through the RZZ pathway, independently of the KBB pathway. We show that hSpindly is a dynamic protein that oligomerizes at unattached kinetochores. Importantly, we identify a specific residue, threonine 552, as critical for hSpindly’s function and mobility. A non-phosphorylatable mutant (T552A) stabilizes hSpindly at kinetochores, impairs SAC signaling, and increases cellular resistance to antimitotic drugs. Altogether, our findings identify hSpindly as a novel dynamic modulator of SAC functionality via the RZZ pathway and highlight it as a potential therapeutic target for overcoming resistance to mitotic inhibitors.

## INTRODUCTION

In human cells, faithful chromosome segregation is ensured by a highly sophisticated surveillance mechanism known as the Spindle Assembly Checkpoint (SAC). SAC activity is primarily regulated by a set of spindle-associated proteins, including Mad1, Mad2, Bub1, BubR1, Bub3, and kinases Mps1 and Aurora B, which are recruited to unattached and/or tensionless kinetochores to initiate checkpoint signalling. The outer kinetochore, a multi-protein structure responsible for tethering chromosomes to spindle microtubules, contains the KMN network (composed of KNL1, Mis12, and Ndc80), which plays a central role in recruiting SAC proteins. KNL1 serves as a scaffold that, upon phosphorylation by Mps1, facilitates the recruitment of Bub1, BubR1, and Bub3. Bub1, in turn, recruits the Mad1-Mad2 complex. Once the SAC is activated, soluble Mad2 (O-Mad2) is converted into its closed conformation (C-Mad2), which binds Cdc20 together with BubR1 and Bub3 to form the mitotic checkpoint complex (MCC), a potent inhibitor of the Anaphase-Promoting Complex/Cyclosome (APC/C)^1,2^. In addition, the kinetochore has an outermost dense layer called fibrous corona that aids microtubule capture and contributes to the initial phases of chromosome alignment in prometaphase. This layer is formed by the Rod-Zw10-Zwilch complex (called RZZ) and the Dynein-dynactin adaptor hSpindly^3,4^. The RZZ acts as another Mad1-Mad2 recruitment to form an additional MCC pool^5^. However, the precise mechanisms by which the RZZ pathway regulates SAC remain poorly understood.

hSpindly is a protein of 605 amino acids with a conserved 32-amino acid motif found in a break between predicted coiled-coil domains in the amino-terminus^6,7^. Interphase localization of hSpindly is mainly nuclear. In mitosis, hSpindly localizes to kinetochores in early prometaphase. After chromosome alignment is achieved, hSpindly leaves the kinetochores and migrates to the spindle poles. At later stages of mitosis (anaphase and telophase), hSpindly is not visible in any spindle structures, being degraded upon mitotic exit^6^. hSpindly undergoes extensive phosphorylation during mitosis, with putative phosphorylation sites targeted by Cdk1 and Mps1 at its carboxy-terminal region, although the functional significance of these modifications remains unclear^8,9^. The recruitment of hSpindly to kinetochores is dependent on the RZZ complex and requires two distinct post-translational modifications: farnesylation at the carboxy-terminus and Mps1-mediated phosphorylation of the amino-terminal domain^10,11^. This phosphorylation triggers conformational changes in hSpindly that promote oligomerization of the RZZ–hSpindly complex, a key step for its proper function during mitosis. This oligomerization is critical for the assembly of the fibrous corona^3^. Moreover, hSpindly acts as a negative regulator of SAC signalling in a manner that is independent of the dynein-mediated stripping pathway^12^. However, the precise mechanism by which hSpindly functions in SAC activation remains unknown.

Importantly, hSpindly is overexpressed in several tumour types associated with poor prognosis and resistance to chemotherapeutic agents, including antimitotic drugs^13,14^. These findings underscore the clinical relevance of understanding the regulatory mechanisms involving hSpindly.

In this study, we identify hSpindly as a novel regulator of SAC activation that operates through the RZZ pathway under the control of Bub1. We demonstrate that hSpindly is dynamically localized to unattached kinetochores, and how its recruitment depends on phosphorylation at threonine 552. A non-phosphorylatable mutant (T552A) disrupts hSpindly dynamics, compromises SAC signalling, and increases resistance to antimitotic agents. These findings reveal a new mechanism by which hSpindlýs dynamic contributes to checkpoint fidelity and therapeutic response.

## RESULTS

### 1. hSpindly interacts with SAC proteins in the absence of microtubules

hSpindly is a cell cycle-regulated protein involved in fibrous corona formation. Its levels increase progressively throughout the cell cycle and it becomes phosphorylated during mitosis^8^ (Supplemental Figure 1A). Given its localization and regulation during mitosis, we hypothesized that hSpindly might participate directly in SAC activation. To explore this possibility, we examined whether hSpindly physically associates with core SAC components under microtubule-disrupting conditions in the presence of high concentration of nocodazole.

To this end, we first generated inducible RPE-1 cells expressing GFP-tagged hSpindly. Western blot analysis confirmed that GFP-hSpindly expression (hSpindly^EXO^) is higher than the endogenous protein (hSpindly^ENDO^) and is strictly dependent on doxycycline induction (Figure 1A). Next, we compared the localization of hSpindly^EXO^ and hSpindly^ENDO^ at the fibrous corona in mitotic cells. To mark the kinetochore, we used CENP-C and Bub1, both well-established kinetochore components. As expected, hSpindly^EXO^ is localized at kinetochores (Figure 1B, Supplemental 1B), displaying a localization pattern consistent with the endogenous protein. Quantitative analysis revealed that hSpindly^EXO^ exhibits a broader distribution compared to the endogenous protein, extending further into the fibrous corona. While Bub1 levels were not altered by hSpindly overexpression, its distribution pattern was slightly modified (Figure 1C, Supplemental Figure 1C). These findings confirm that hSpindly is overexpressed and correctly localized at kinetochores.

**Figure 1.**
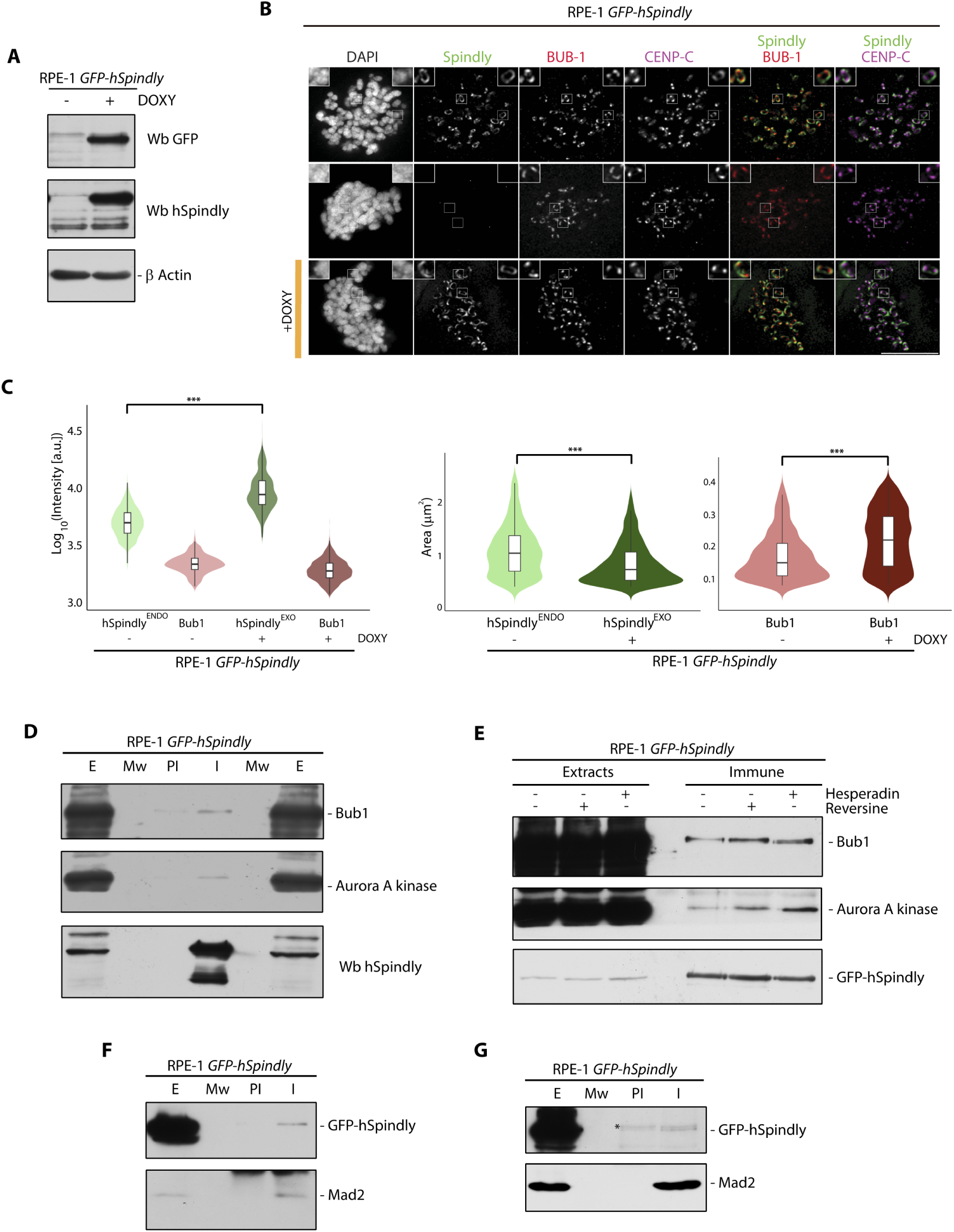
hSpindly as a component of SAC machinery. **A.** Stable inducible RPE-1 *GFP-hSpindly* cells were treated or not with doxycycline (DOXY). Cell extracts were prepared and immunoblotted with the indicated antibodies. In Wb hSpindly, GFP-hSpindly is the upper band and hSpindly endogenous is the lower band. **B.** Representative images of hSpindly levels at kinetochores of RPE-1 *GFP-hSpindly* treated or not with DOXY in the presence of 3.3µM nocodazole (NZ). Endogenous or exogenous hSpindly were visualized by hSpindly antibody or GFP, respectively. Bub1 or CENP-C antibodies were used as kinetochore markers, and DAPI to visualize the DNA. Scale bar: 10μm; insets, 2x zoom. **C.** Violin plots representing intensity or total area of hSpindly^ENDO^, hSpindly^EXO^ and Bub1 at kinetochores from the experiment shown in panel B. Lines indicate mean or standard deviation. Statistical analysis was performed with a non-parametric t-test comparing two unpaired groups (***, p<0.001). **D.** RPE-1 *GFP-hSpindly* cell extracts after NZ and DOXY treatment for 24 hours were immunoprecipitated with anti-hSpindly (I) or normal mouse serum (PI) as control. Immunoprecipitated materials were analyzed by western blot using the antibodies indicated. **E.** Stable inducible RPE-1 *GFP-hSpindly* cells were treated with DOXY and NZ for 24 hours. Before collecting, mitotic cells were incubated with 0.5μM hesperadin or 1μM reversine in the presence of 20μM proteasome inhibitor MG132 for 90 minutes. Cell extracts were used to immunoprecipitate as in D. **F, G** RPE-1 *GFP-hSpindly* cell extracts treated with NZ and DOXY for 24 hours were immunoprecipitated using anti-Mad1 or anti-Mad2 antibodies (I), with normal mouse serum used as a control (PI). Anti-Mad2 was used as a control of Mad1 immunoprecipitation. Western blot analysis was performed using the specified antibodies. E: Extracts; Mw: Molecular weight; I: Immune; PI: Preimmune. The upper band marked with asterisk is an unspecific band.

To determine whether hSpindly contributes to SAC activation independently of the dynein-mediated stripping pathway, we performed all subsequent experiments in the presence of high doses of nocodazole (NZ) to depolymerize microtubules. We then assessed whether hSpindly forms complexes with SAC proteins by immunoprecipitation followed by western blotting. As shown in Figure 1D, hSpindly co-immunoprecipitated Bub1 and Aurora A kinase. These interactions persisted in the presence of hesperadin (an Aurora B inhibitor) and reversine (an Mps1 inhibitor), indicating that hSpindly binding to Bub1 and Aurora A is independent of Mps1 and Aurora B activity (Figure 1E). Given the central role of Mad1–Mad2 recruitment at kinetochores in SAC signalling, we also tested whether hSpindly associates with these components. Indeed, hSpindly formed immunocomplexes with both Mad1 and Mad2 (Figure 1F and 1G), further supporting the notion that hSpindly engages with key SAC regulators and may directly contribute to SAC activation.

### 2. Overexpression of hSpindly causes a delay in mitotic exit

To investigate the functional consequences of hSpindly overexpression, we focused on mitotic exit, a process tightly regulated by SAC silencing. Once SAC is inactivated, the level of Cyclin B and Securin is reduced since both are degraded by the APC/C^Cdc20^ proteasome, allowing progression into anaphase^15,16^. Therefore, monitoring Cyclin B levels serves as a functional readout of SAC inactivation under conditions of hSpindly overexpression, employing two complementary experimental strategies, one that allows microtubule regrowth and another that continues in the presence of microtubule disruption. In the first approach, cells were treated with NZ for 24 hours to depolymerize microtubules and fully activate the SAC. Afterward, NZ was washed out to allow microtubule reformation, and cells were harvested at 30-minute intervals. Western blot analysis revealed that Cyclin B degradation was delayed in cells overexpressing hSpindly compared to controls expressing only endogenous levels, indicating a delay in mitotic exit (Figure 2A).

**Figure 2.**
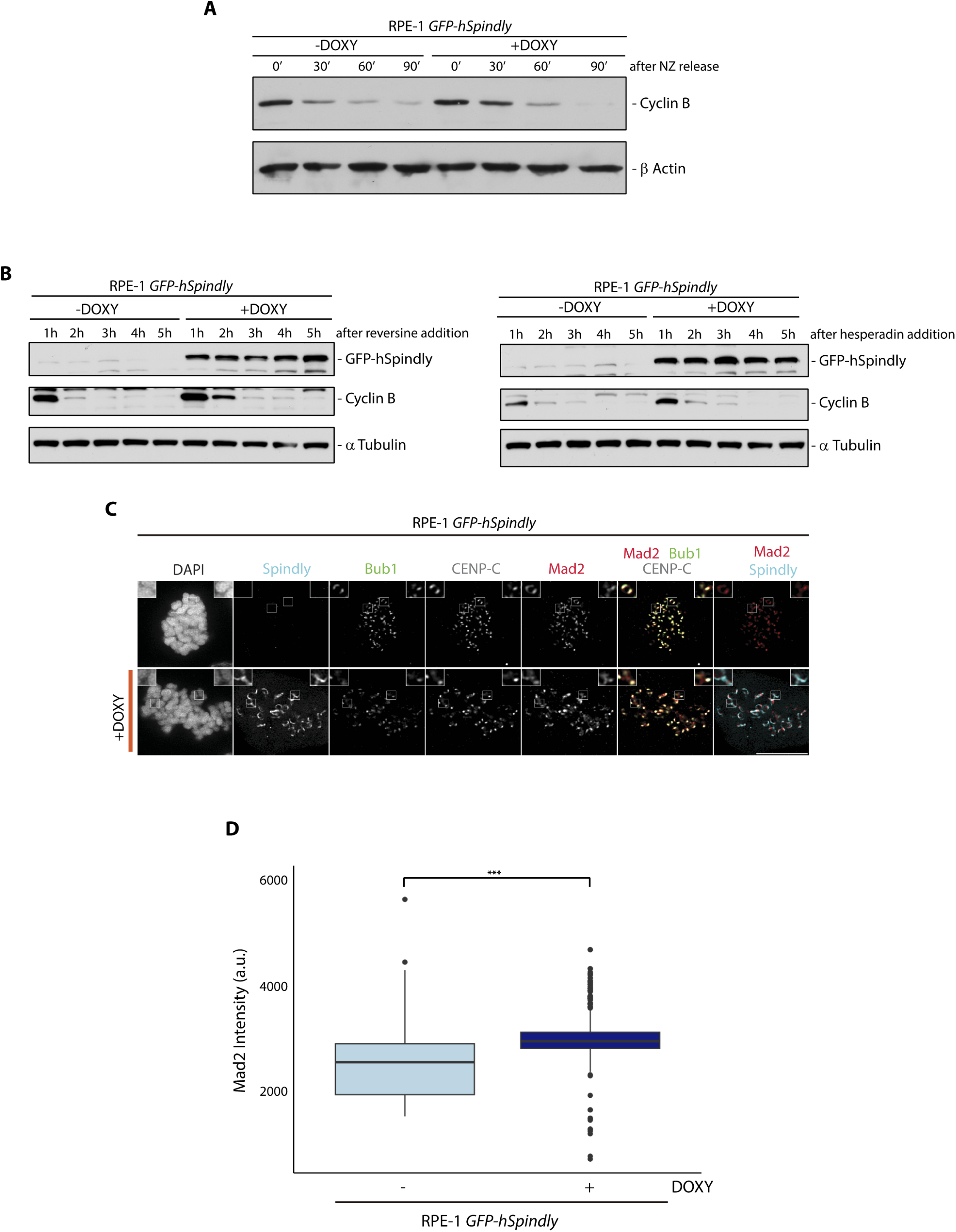
hSpindly overexpression maintains SAC activation by enhancing Mad2 recruitment at kinetochores. **A.** RPE-1 *GFP-hSpindly* cells were washed out after 24 hours of NZ treatment without or with DOXY and collected every 30 minutes. Cell extracts were analyzed by western blot with the antibodies indicated. **B.** RPE-1 *GFP-hSpindly* cells were treated with NZ with or without DOXY for 24 hours and collected after 1μm reversine (left panel) or 0.5μm hesperadin (right panel) addition at indicated times. Cell extracts were prepared and immunoblotted with the indicated antibodies. **C.** Representative images of GFP-hSpindly, Bub1, CENP-C or Mad2 levels at kinetochores in RPE-1 *GFP-hSpindly* cells treated or not with DOXY in the presence of NZ. DAPI to visualize the DNA. Scale bar: 10μm; insets, 2x zoom. **D.** Box plots representing intensity of Mad2 at kinetochores from the experiment shown in panel C. Black lines indicate mean. Statistical analysis was performed with a nonparametric t-test comparing two unpaired groups (***, p<0.001).

The second approach aimed to assess SAC silencing in the continued absence of microtubules by chemically inactivating SAC kinases. Mps1 and Aurora B are essential for both SAC establishment and maintenance, and their inhibition using reversine and hesperadin, respectively, can override the checkpoint and drive mitotic exit even in the presence of unattached kinetochores^17,18^. HCT116 and RPE-1 cells were treated with NZ and subsequently exposed to reversine or hesperadin for 90 minutes (Supplemental Figure 2A). Reversine alone effectively reduced Cyclin B levels in both cell types, whereas hesperadin only produced this effect in HCT116 cells after this time of treatment. In RPE-1 cells, Cyclin B levels remained high in the presence of hesperadin. The use of the proteasome inhibitor MG132 confirmed that Cyclin B degradation was proteasome-dependent. Additionally, differences in BubR1 phosphorylation patterns between the two cell lines were observed following kinase inhibition, suggesting that SAC robustness and regulation may differ between RPE-1 and HCT116 cells^19^. Prolonged reversine or hesperadin treatment of RPE-1 cells overexpressing hSpindly (Figure 2B) revealed that Cyclin B levels remain elevated after the addition of the inhibitors. This reinforces the conclusion that hSpindly overexpression impairs SAC silencing and delays mitotic exit. To gain mechanistic insight into this delay, we analysed the kinetochore localization of Mad2 in cells with unattached kinetochores. Fluorescence microscopy revealed increased Mad2 intensity at kinetochores in cells overexpressing hSpindly (Figure 2C, Figure 2D, Supplemental Figure 2B). Hence, hSpindly contributes to regulate the presence of Mad2 at kinetochores.

### 3. Mad2 is recruited at kinetochores via hSpindly independently of Bub1, regulating the effectiveness of anti-mitotic drugs

Previous studies have demonstrated an interplay between Bub1 and the RZZ complex in maintaining SAC functionality^20,21^. Bub1 is a serine/threonine kinase, and its catalytic activity plays a minor role in SAC activation^22^. However, given that hSpindly interacts with Bub1, and that Bub1 phosphorylates hSpindly at least *in vitro*^4^, we asked whether inhibition of Bub1 kinase activity might affect the interaction between hSpindly and the Mad1/Mad2 complex (Figure 1). To explore this, we used BAY 1816032, a specific inhibitor of Bub1 kinase activity^23,24^. Strikingly, immunoprecipitation of hSpindly under Bub1-inhibited conditions showed increased association with Mad1, while Bub1 binding was reduced (Figure 3A). These results suggest that Bub1 kinase activity negatively regulates the interaction between hSpindly and the Mad1/Mad2 complex.

**Figure 3.**
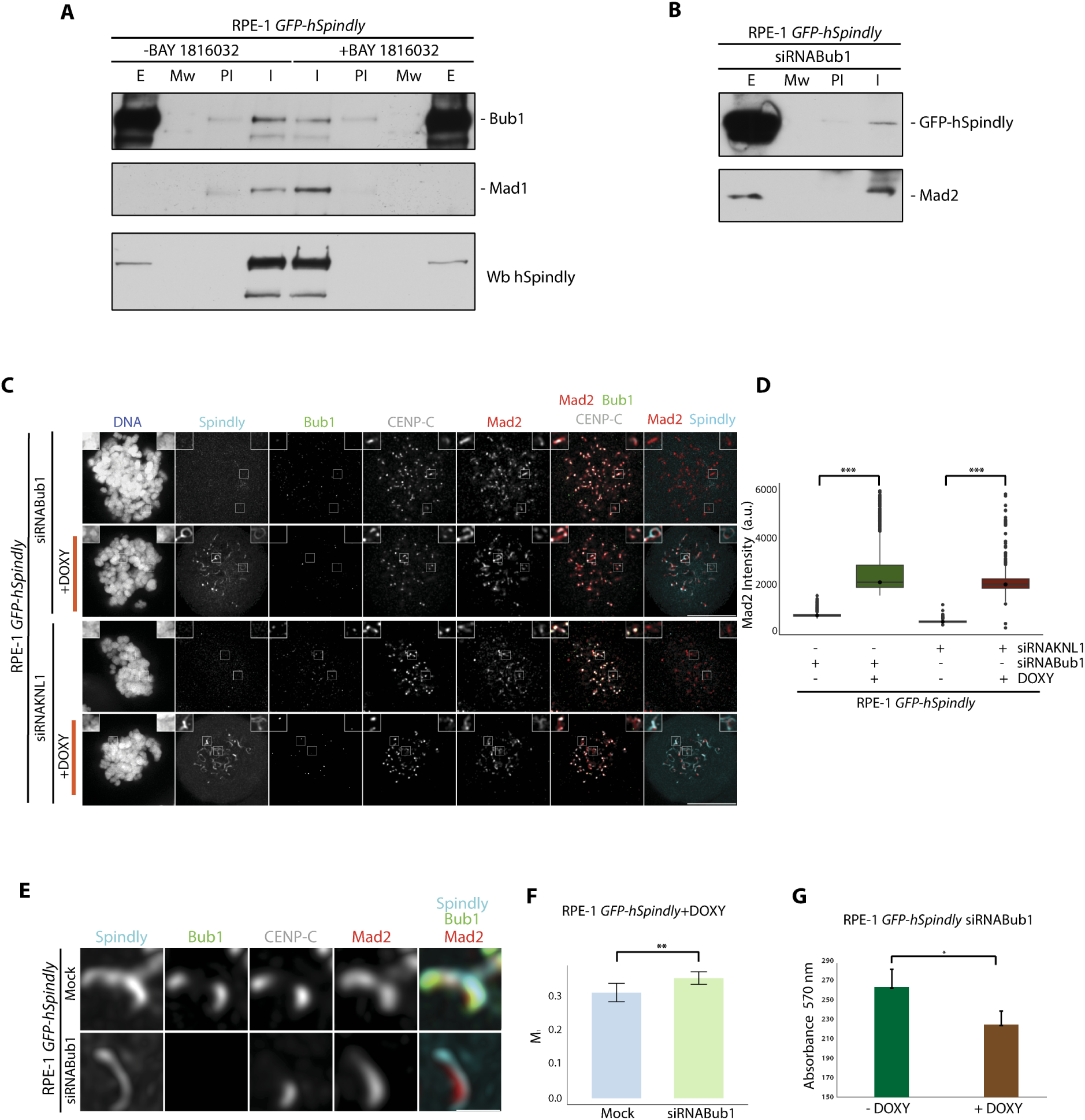
Mad1-Mad2 complexes are recruited at kinetochores in the absence of the KBB pathway. **A.** RPE-1 *GFP-hSpindly* prometaphase cells induced by DOXY were treated or not with 10μM BAY 1816032 inhibitor for 5 hours in the presence of 20μM MG132. Cell extracts were used to immunoprecipitate hSpindly (I) and normal mouse serum (PI) as a control. Immunoprecipated material was analyzed by western blot. **B.** RPE-1 *GFP-hSpindly* were transfected with siRNABub1 for 48 hours. Cells were treated with NZ and DOXY 24 hours before harvesting. Extracts were used to immunoprecipitate Mad1 (I) using normal mouse serum as a control (PI). Complexes were analyzed by immunoblotting with antibodies as indicated. **C.** Representative images of RPE-1 *GFP-hSpindly* prometaphase cells in the presence or not of DOXY after Bub1 or KNL1 siRNA treatment for 48 hours. Scale bar: 10μm; insets, 2x zoom. **D.** Scatter dot boxes representing intensity of Mad2 at kinetochores from the experiment shown in panel C. Black lines indicate mean. Statistical analysis was performed with a nonparametric t-test comparing two unpaired groups. (***, p<0.001). **E.** Representative images of a magnification of kinetochores in RPE-1 *GFP-hSpindly* prometaphase cells after 48 hours of Bub1 siRNA in the presence of DOXY. Scale bar: 10μm. **F.** Box plots representing colocalization of Mad2-hSpindly at kinetochores from the experiment shown in panel E. The Manders’ M_1_ coefficient is shown. Statistical analysis was performed with a parametric model comparing two unpaired groups. (**, p<0.01). **G.** Colony assays showing cell viability for RPE-1 *GFP-hSpindly* cells transfected with siRNABub1 were performed by measuring absorbance 570nm and represented in graphs. Error bars represent the SD (n=3). *p<0.05 (Student’s t-test). E: Extracts; Mw: Molecular weight; I: Immune; PI: Preimmune.

Mad1/Mad2 recruitment to unattached kinetochores is known to occur through two parallel pathways: the KNL1-Bub3-Bub1 (KBB) pathway and the RZZ pathway. Since Bub1 acts as a scaffolding protein to recruit Mad1/Mad2 via the KBB pathway, we next asked whether hSpindly could recruit Mad1/Mad2 by itself serving as a key mediator in the RZZ pathway. To test this, we performed an RNAi-mediated knockdown of Bub1 in RPE-1 GFP-hSpindly induced cells treated with NZ. Surprisingly, even upon Bub1 depletion, hSpindly maintained its interaction with Mad1 (Figure 3B, Supplemental Figure 3A).

To functionally assess whether hSpindly can support Mad2 localization in the absence of Bub1, we overexpressed *GFP-hSpindly* in Bub1-depleted cells. As expected, Bub1 knockdown caused a marked reduction in Mad2 kinetochore levels; however, this reduction was rescued by *hSpindly* overexpression (Figure 3C, Figure 3D, Supplemental Figure 3A). A similar rescue was observed following depletion of KNL1, a core component of the KBB pathway. Mad2 levels at kinetochores were reduced upon KNL1 knockdown but were restored upon doxycycline-induced hSpindly expression (Figure 3C, Figure 3D, Supplemental Figure 3B). Notably, in Bub1-depleted cells, Mad2 localization shifted and was found to colocalize with hSpindly around the kinetochore, suggesting that hSpindly not only promotes Mad2 recruitment but also influences its spatial distribution (Figure 3E, Figure 3F). Collectively, these data support a model in which hSpindly recruits Mad2 to unattached kinetochores independently of the KBB pathway and helps regulate its distribution within the fibrous corona.

One strategy used in cancer treatment is manipulation of spindle dynamics. By (de)stabilising the microtubules, anti-mitotic drugs prolong the SAC and thus block the metaphase-anaphase transition. This result in a long-term SAC-dependent mitotic arrest followed by cell death, compromising tumor cell proliferation^25^.However, many tumour cells have a defective SAC which does not respond to microtubules-interfering drugs, reducing the efficiency of the treatment^26^. Alterations in the expression levels of SAC components have been linked to tumour progression and resistance to anti-mitotic therapies. For instance, proteins such as Mad2, BubR1, Aurora B, and Mps1 are frequently overexpressed in various types of cancer^27–29^. Given that we have demonstrated a role for hSpindly in SAC regulation, we next asked whether hSpindly overexpression would increase the sensitivity to NZ-induced microtubule destabilization in the absence of Bub1. To investigate the response to antimitotic drugs, we performed a cell viability assay. We compared the number of colonies that appeared after 24 hours in the presence of NZ in Bub1 knockdown cells with or without hSpindly overexpression. As shown in Figure 3G, hSpindly overexpression increases the sensitivity of cells to NZ, reducing the cell viability. This could potentially be the mechanism underlying the reduced effectiveness of anti-mitotic drugs in tumour cells with an altered expression of SAC proteins.

### 4. The phosphorylation state of hSpindly affects the recruitment of Mad2 at unattached kinetochores

Given that hSpindly plays a key role in SAC activation, understanding how this protein is regulated within this network is of particular interest. hSpindly is phosphorylated during mitosis, as indicated by a mobility shift in SDS-PAGE upon NZ treatment of RPE-1 cells. This shift was enhanced by okadaic acid (a phosphatase inhibitor) and abolished by λ-phosphatase treatment (Supplemental Figure 4A). Previous studies identified putative CDK1 and Mps1 phosphorylation sites within the carboxy-terminal region of hSpindly^8,9^, although the functional relevance of these sites remains largely unknown.

In previous works from our lab, we generated a phospho-specific antibody against hSpindly phosphorylated at threonine 552, which specifically recognized mitotic cells (unpublished data). To investigate the functional role of this modification, we generated an inducible RPE-1 cell line expressing a non-phosphorylatable mutant form of hSpindly (T552A) tagged with GFP (hSpindly-T552A^EXO^) (Supplemental Figure 4B). Western blot analysis confirmed doxycycline-dependent expression of hSpindly-T552A^EXO^, albeit at lower levels compared to wild-type hSpindly^EXO^. This suggests that phosphorylation at T552 may contribute to protein stability (Figure 4A).

**Figure 4.**
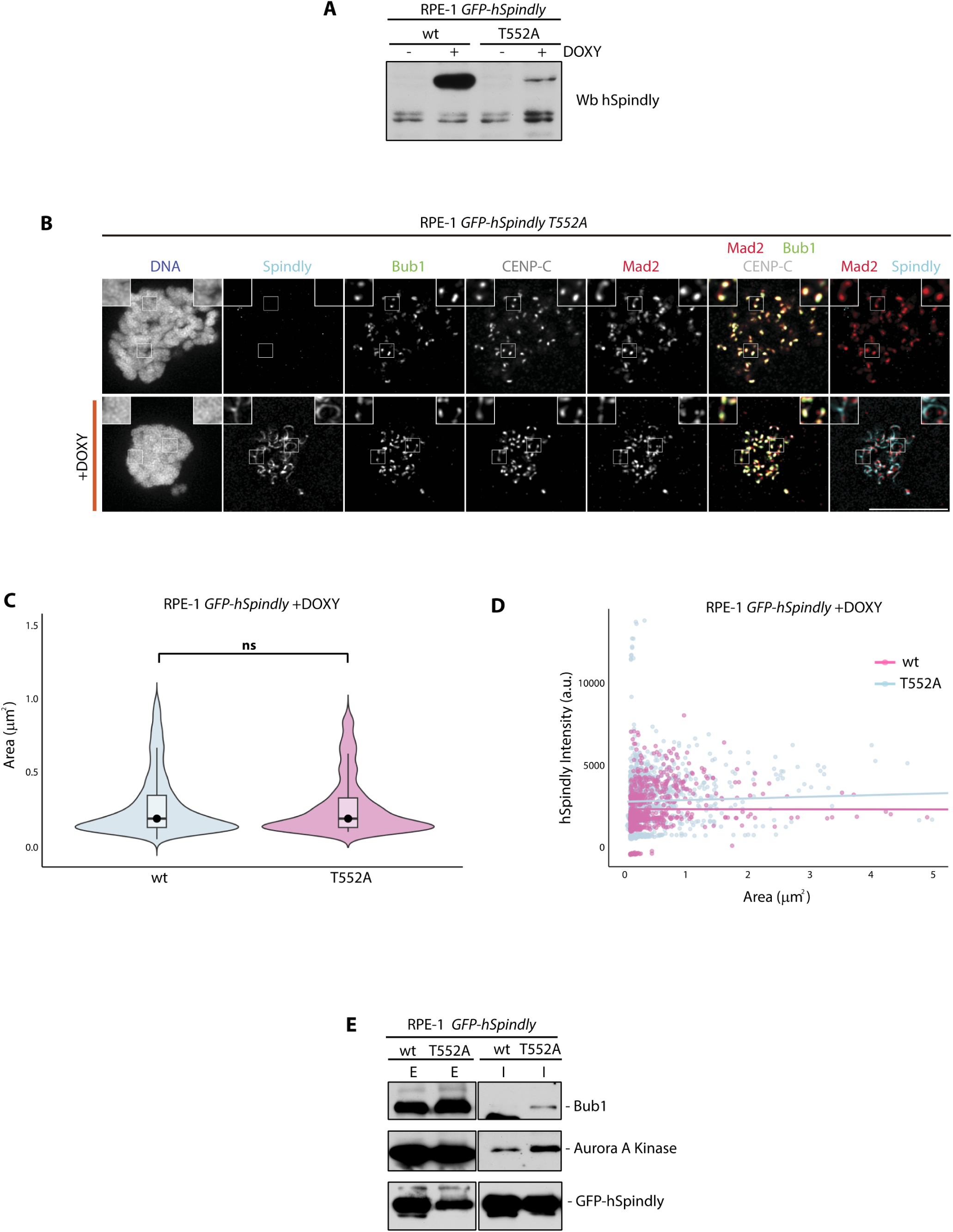
hSpindly T552A is localized at kinetochore as wild-type and interacts with SAC proteins. **A.** Stable inducible RPE-1 *GFP-hSpindly wild-type* (wt) or *GFP-hSpindly T552A* (T552A) cells were treated or not with doxycycline (DOXY) in the presence of NZ for 24 hours. Cell extracts were prepared and immunoblotted with the indicated antibodies. Endogenous hSpindly (lower band) was used as loading control **B.** Representative images of hSpindly T552A levels at kinetochores of RPE-1 *GFP-hSpindlyT552A* prometaphase cells treated or not with DOXY. hSpindlyT552A or Mad2 were visualized by GFP or anti-Mad2, respectively. Bub1 or CENP-C antibodies were used as kinetochore markers, and DAPI to visualize the DNA. Scale bar: 10μm; insets, 2x zoom. **C.** Violin plots representing intensity or total area of hSpindly T552A compared with hSpindly wild-type (data Figure 1C). Black dots or lines indicate mean or standard deviation. Statistical analysis was performed with a nonparametric t-test comparing two unpaired groups. (ns, not significant). **D.** Distribution of hSpindly^EXO^ or hSpindly-T552A^EXO^ populations based on kinetochore signal intensity and area. Each dot represents a kinetochore; lines indicate the mean. **E.** RPE-1 *GFP-hSpindly wt* or *GFP-hSpindly T552A* induced cell extracts after NZ treatment were immunoprecipitated with GFP-Trap beads. Immunoprecipitated materials were analyzed by western blot using the antibodies indicated. E: Extracts; I: Immune.

Microscopy analysis showed that hSpindly-T552A^EXO^ was correctly targeted to kinetochores, as indicated by colocalization with the kinetochore markers Bub1 and CENP-C in prometaphase cells (Figure 4B, Supplemental 4C). Its localization pattern and intensity at the fibrous corona were comparable to those of wild-type hSpindly^EXO^ (Figure 4C, Figure 4D). However, immunoprecipitation experiments revealed that the T552A mutant bound increased amounts of Bub1 and Aurora A kinase relative to the wild-type protein (Figure 4E). To determine whether phosphorylation at T552 affects Mad2 recruitment to unattached kinetochores, we examined Mad2 localization in the context of Bub1 depletion and overexpression of hSpindly-T552A^EXO^ (Figure 5A, Figure 5B). While hSpindly^EXO^ (Figure 3C) rescued Mad2 recruitment in Bub1-deficient cells, hSpindly-T552A^EXO^ failed to do so. Furthermore, no enhanced colocalization between Mad2 and hSpindly-T552A was observed around kinetochores. In this case, Mad2 distribution resembled that observed with Bub1 depletion (Figure 5C, Figure 5D).

**Figure 5.**
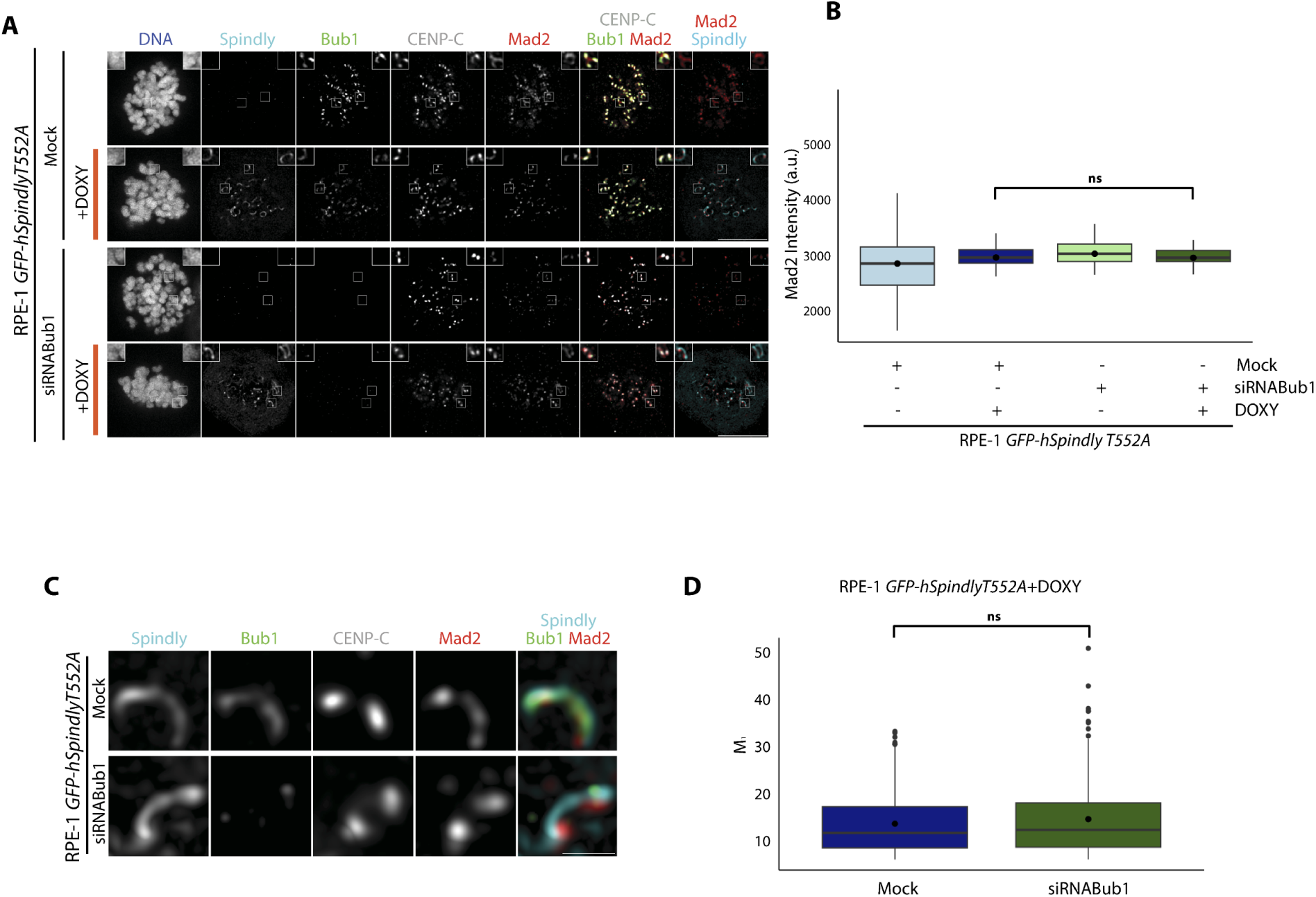
hSpindly T552A recruits Mad2 at kinetochores independently of Bub1. **A.** Representative images of *RPE-1 GFP-hSpindlyT552A* prometaphase cells in the presence or not of DOXY after Mock or Bub1 siRNA treatment for 48 hours. GFP-hSpindly T552A, Bub1, CENP-C, Mad2 were visualized with green fluorescence protein or by using the indicated antibodies. Scale bar: 10μm; insets, 2x zoom. **B.** Box plots representing intensity of Mad2 at kinetochores from the experiment shown in panel A. Statistical analysis was performed with a nonparametric t-test comparing two unpaired groups. **C.** Representative images of a magnification of Kinetochores in RPE-1 *GFP-hSpindly T552A* prometaphase cells after 48 hours of Mock or Bub1 siRNA in the presence of DOXY. Scale bar: 10μm. **D.** Box plots representing colocalization of Mad2-hSpindly at kinetochores from the experiment shown in panel C. The Manders’ M_1_ coefficient is shown. Black lines indicate mean. Statistical analysis was performed with a nonparametric t-test comparing two unpaired groups. (ns, not significant).

Together, these results indicate that phosphorylation of hSpindly at threonine 552 is required for its ability to promote Mad2 recruitment to unattached kinetochores, particularly in the absence of Bub1. This phosphorylation likely plays a key role in coordinating SAC signalling via the RZZ pathway.

### 5. hSpindlýs dynamic behaviour depends on the threonine 552 phosphorylation and Bub1

hSpindly localizes to the fibrous corona through its interaction with the RZZ complex, with which it oligomerizes *in vitro*. Structurally, hSpindly forms a dimer composed of four interacting coiled-coil segments^3,4,11^. However, its dynamic behaviour *in vivo* remains poorly understood. To investigate hSpindly mobility at unattached kinetochores, we applied live cell imaging in combination with Raster Image Correlation Spectroscopy (RICS), a quantitative fluorescence microscopy technique that extracts diffusion coefficients and binding-unbinding constants from spatial and temporal fluorescence fluctuations during confocal scanning^30^.

RPE-1 cells expressing hSpindly^EXO^ or hSpindly-T552A^EXO^ were treated with NZ to generate unattached kinetochores. RICS measurements were performed both at kinetochores and in the cytoplasm, subtracting the immobile fraction every 4 frames, and modelled respectively as a binding model (green box) and a diffusion model (red box) of Supplemental Figure 5. hSpindly^EXO^ exhibited dynamic behaviour consistent with transient binding at unattached kinetochores and a diffusive movement away from the kinetochores. In the same manner, hSpindly-T552A^EXO^ displayed a similar time constant (directly proportional to the on and off binding times). Upon Bub1 depletion, no difference in these values was observed. Given that hSpindly seems to be moved driven by a binding mechanism in the kinetochores, a more exhaustive RICS analysis was performed in that area. Evaluation of the binding/unbinding mechanism to fixed locations was performed by subtracting the immobile fraction after 4 and 10 frames and the shape of the spatial correlation function was evaluated. Figure 6A displays the spatial correlation function of hSpindly^EXO^, hSpindly-T552A^EXO^ and hSpindly^EXO^ upon the depletion of Bub1 (from left to right), subtracting the immobile fraction every 4 (upper) or 10 (bottom) frames. hSpindly^EXO^ clearly shows a change in the shape of the correlation function, suggesting there is a transitory binding process. In contrast, hSpindly-T552A^EXO^ and hSpindly^EXO^ upon the depletion of Bub1 do not show a clear difference in the shape of the correlation function suggesting the binding to fixed locations. In order to quantify this effect, correlation functions were fitted and time constant (⍰) was subtracted (Figure 6A, graph). Clearly wild-type protein displays a transitory binding mechanism with respect to hSpindly-T552A^EXO^ or hSpindly^EXO^ upon Bub1 depletion that bind to fixed locations.

**Figure 6.**
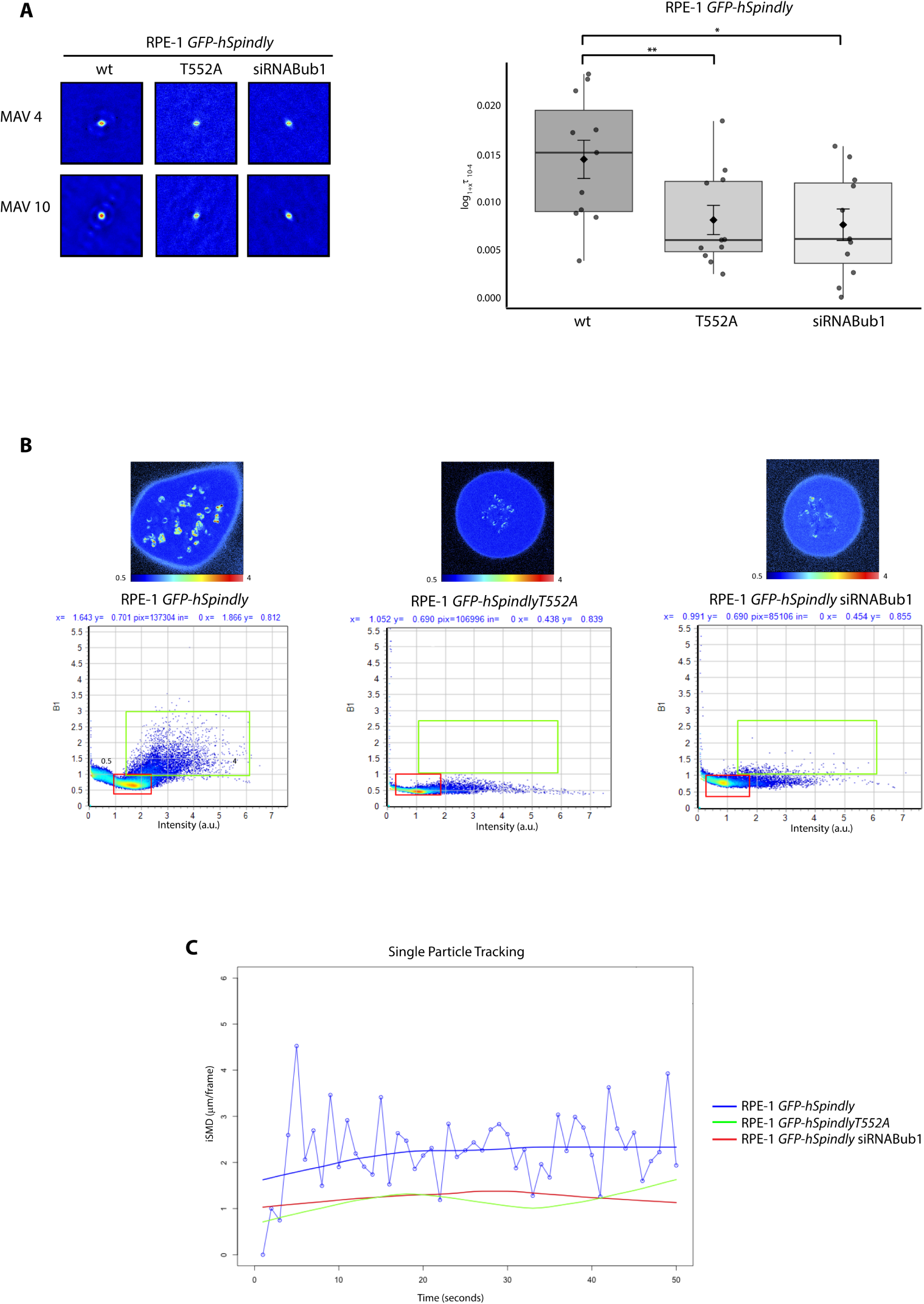
hSpindly is a dynamic protein at kinetochores. **A.** Spatial correlation functions of hSpindly^EXO^, hSpindly-T552^EXO^ and hSpindly^EXO^ upon Bub1 knockdown after the subtraction of the immobile fraction every 4 and 10 frames (left) and the graph representing the difference of the time constant when the correlation functions are fitted after the immobile fraction removal every 4 and 10 frames (right). (*, p<0.05; **, p<0.01). **B.** 512x512 Brightness micrographs with their respective Brightness/Intensity plot of hSpindly, hSpindly-T552 and hSpindly upon Bub1 depletion. Image size 25.56µmx25.56µm. **C.** Single Particle Tracking analysis (SPT) on kinetochores of hSpindly, hSpindly-T552 and hSpindly upon Bub1 depletion.

To further investigate the mechanistic basis of these effects, we used the Number and Brightness (N&B) method to assess hSpindly oligomerization in live cells. N&B is a fluorescence-based approach that measures pixel-by-pixel intensity fluctuations to infer protein aggregation states, allowing the distinction between monomeric and oligomeric forms in living cells. In this context, when plotting brightness versus fluorescence intensity, oligomeric forms will show an increase in both brightness and intensity. In contrast, aggregated forms are considered immobile, so an increment brightness is not detected in those pixels, while fluorescence intensity appears highly increased. Following NZ treatment, we compared the oligomerization profiles of hSpindly^EXO^,hSpindly-T552A^EXO^, and hSpindly^EXO^ under Bub1 knockdown conditions. As shown in Figure 6B, hSpindly displayed increasing brightness values, consistent with a higher oligomerization state. In contrast, both hSpindly-T552A^EXO^ and hSpindly^EXO^ under Bub1 depletion showed significantly lower brightness values, while the intensity profile remained unchanged, indicating a higher aggregation state but reduced mobility at the kinetochores. It is noteworthy that the difference in this analysis may be due to the movement of kinetochores. As shown also in Figure 6A, hSpindly binds/unbinds in an equilibrium, due to the NZ treatment, and this binding mechanism may infer the difference of the movement of the kinetochores. For this reason, Single Particle Tracking (SPT) analysis was performed on the whole kinetochores, demonstrating faster kinetochore dynamics in the wild-type sample (Figure 6C and Supplemental movies 1 to 3). Therefore, hSpindly-T552A^EXO^ is aggregated in the kinetochores, allowing binding but preventing unbinding, which suggests that the self-assembly mechanism lacks equilibrium in the mutated protein. This generates that, upon T552A mutation, hSpindly remains more tightly attached to the kinetochore, making these structures more resistant to disassembly. Interestingly, depletion of Bub1 generates a similar dynamic behaviour.

### 6. Resistance to antimitotic drugs is enhanced upon loss of threonine 552 phosphorylation

Given that threonine 552 phosphorylation regulates hSpindly’s role in recruiting SAC proteins to unattached kinetochores, we next examined whether this post-translational modification influences cellular sensitivity to antimitotic agents. To this end, we performed colony formation assays under doxycycline-induced overexpression of either hSpindly^EXO^ or hSpindly-T552A^EXO^, in cells depleted of hSpindly^ENDO^ and KNL1. Cells were exposed to NZ for 24 hours to induce mitotic arrest, followed by washout and recovery for two weeks. Colony formation was assessed via crystal violet staining. Notably, overexpression of hSpindly-T552A^EXO^ resulted in a significantly higher number of surviving colonies compared to hSpindly^EXO^, indicating that the loss of phosphorylation at threonine 552 promotes resistance to antimitotic treatment (Figure 7A). These findings underscore the functional relevance of threonine 552 phosphorylation in regulating SAC robustness and drug sensitivity.

**Figure 7.**
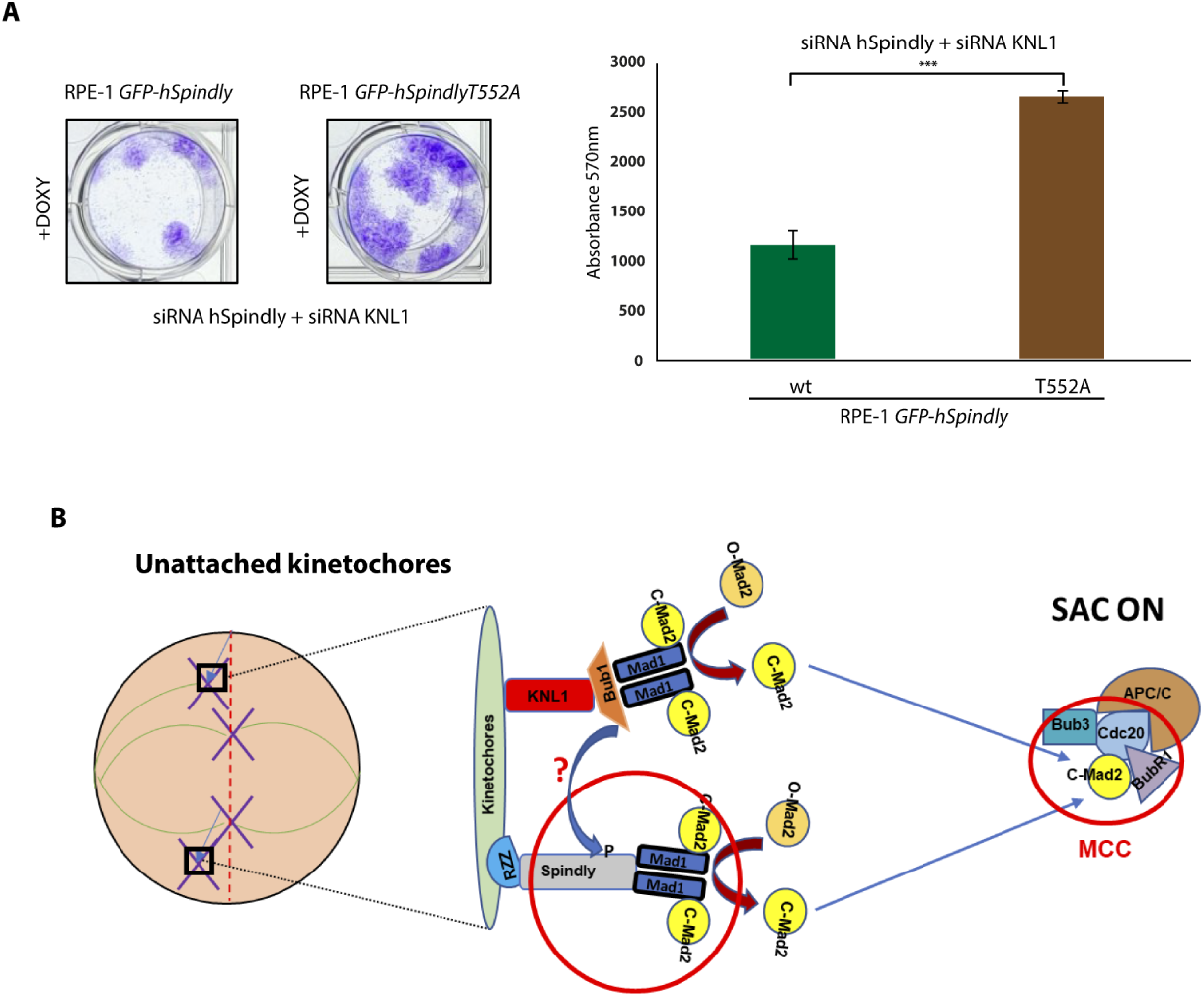
Phosphorylation of threonine 552 determines cellular resistance to anti-mitotic agents. **A.** Colony assays showing cell viability of RPE-1 *GFP-hSpindly* or RPE-1 *GFP-hSpindly T552A* were performed after 24 hours of NZ treatment in the presence of DOXY and in the absence of both endogenous hSpindly and KNL1. Colony forming was quantified by measuring absorbance 570nm and represented in graphs. Error bars represent the SD (n=3). ***, p<0.001 (Student’s t-test). **B.** Model to illustrate how spindle checkpoint regulation is controlled by the KBB and RZZ pathways at unattached kinetochores. Bub1 kinase activity likely controls the activation of the RZZ pathway by the phosphorylation of hSpindly. hSpindly acts as a recruiter of the Mad1-Mad2 complex controlling the MCC formation with the KBB pathway.

Collectively, our data support a novel SAC regulation model in which hSpindly is finely controlled. Notably, phosphorylation of hSpindly at a single residue is sufficient to alter its functionality and dynamism, thereby impacting mitotic progression and cellular response to chemotherapy.

## DISCUSSION

The fidelity of the SAC is essential for maintaining genome stability. Recent studies have implicated hSpindly in SAC activity. In lower organisms like *C. elegans*, the interaction between MDF-1/Mad1 and SPDL-1/hSpindly has been established^31^. In human cells, proteomics approaches have identified hSpindly within the Mad1 proximity network in both HeLa and RPE-1 cells^23^. Moreover, point mutants (S256A and F258A) of hSpindly that prevent dynein recruitment can lead to metaphase arrest and cell death, suggesting a more complex role for hSpindly beyond its function as a dynein adaptor^12^. Further investigations have shown that the fibrous corona, composed of the RZZ complex and hSpindly, enhances SAC activation. Specifically, this outermost kinetochore structure facilitates the recruitment and stabilization of Mad1-Mad2 complexes^5,11^. Additionally, Cyclin B1 has been proposed to contribute to Mad1 recruitment at the fibrous corona, further potentiating SAC signalling^32^. These findings underscore the regulatory capacity of the corona in checkpoint signalling and set the stage to explore hSpindly’s role beyond dynein recruitment.

Our results provide further evidence for a direct role of hSpindly in SAC activation. We show that hSpindly interacts with Bub1 and the Mad1–Mad2 complex, and that its overexpression delays degradation of Cyclin B consistent with delayed SAC silencing by Mad1-Mad2 complex accumulation at unattached kinetochores. Previous studies have shown that persistent localization of the Mad1-Mad2 complex can sustain SAC signalling even in cells with properly aligned chromosomes^33^. Remarkably, hSpindly overexpression restores Mad2 localization at unattached kinetochores even in the absence of Bub1 or KNL1, highlighting a Bub1-independent function for hSpindly in SAC activation. hSpindly is recruited to kinetochores through the RZZ complex^4,34^, and our data suggest that its role extends beyond structural anchoring, contributing actively to checkpoint signalling. This supports a model in which hSpindly participates in an RZZ-dependent checkpoint pathway that complements the canonical KNL1-Bub3-Bub1 (KBB) axis. Previous studies positioned RZZ as a recruiter of Mad1-Mad2 to facilitate Bub1-dependent SAC signalling^20,21^. However, our findings reveal a more active role for hSpindly in SAC signalling. Notably, inhibition of Bub1 kinase activity enhances the hSpindly-Mad1 interaction, while Bub1 depletion disrupts hSpindly oligomerization and dynamics at kinetochores. Moreover, the loss of phosphorylation at threonine 552 mimics the effects of Bub1 depletion, reinforcing the notion that Bub1 regulates hSpindly via phosphorylation. Given that Bub1 phosphorylates hSpindly^4^, we propose a model in which Bub1 functions not only through KNL1 scaffolding but also by directly modulating hSpindly activity, thereby bridging the canonical and RZZ-mediated SAC pathways (Figure 7B).

On the other hand, overexpression of hSpindly increases cellular sensitivity to antimitotic drugs, whereas expression of the T552A mutant enhances resistance. This effect becomes particularly evident when the canonical KBB pathway is silenced via depletion of either Bub1 or KNL1. In these experiments, NZ was removed to allow kinetochore-microtubule attachment and subsequent cell cycle progression. Under such conditions, the observed phenotype may reflect hSpindly’s dual role, both in SAC activation and as a key adaptor for dynein–dynactin recruitment at the corona^12,34,35^. hSpindly is recruited to kinetochores through the RZZ complex^4,34^, where it undergoes conformational changes that facilitate dynein binding and activation^36^. These structural transitions depend on oligomerization of the RZZ–hSpindly complex, which is regulated by post-translational modifications, including phosphorylation. Our findings show that the T552A mutation disrupts hSpindly oligomerization and renders it less dynamic at kinetochores. This altered dynamic behaviour, detected by live-cell imaging and biophysical approaches, may prevent the establishment of tension across sister kinetochores, thereby interfering with SAC silencing and timely anaphase onset^5,11^.

Together, our results suggest hSpindly acts as a molecular bridge between the KBB and RZZ pathways being regulated by Bub1 kinase activity. The loss of threonine 552 phosphorylation impairs both checkpoint signalling and mechanical coordination at kinetochores, ultimately contributing to mitotic slippage and resistance to antimitotic therapies. Understanding how SAC signalling and kinetochore mechanics are co-regulated by hSpindly could provide new insights into the fidelity of chromosome segregation and potential mechanisms underlying drug resistance in therapy-refractory cancer cells.

## METHODS

### Cell culture, transfections and drugs

Routinely, RPE-1 and HCT116 cells (from ATCC), and their derivatives were grown in Dulbecco’s Modified Eagle Medium (DMEM) from BioWest (Nuaillé, France) supplemented with 10% heat-inactivated fetal calf serum (Gibco) and 2 mM L-glutamine, 100 U of penicillin/mL, and 100 μg of streptomycin/ml (Gibco) in a 5% CO_2_ humified atmosphere at 37 °C^37^. To reduce gene expression, cells were transfected with siRNA hSpindly (5’ GAA AGG GUC UCA AAC UGA A 3’), siRNA Bub1 (5’ GAG UGA UCA CGA UUU CUA A 3’) or siRNA KNL1 (5’ GGA AUC CAA UGC UUU GAG A 3’) using the Xfect RNA transfection reagent (Takara Bio Inc., Shiga, Japan) following the manufacturer’s standard instructions. In the indicated experiments, cells were treated with nocodazole (3.3μM, Sigma); MG132 (20μM, Santa Cruz Biotechnology); BAY 1816032 (10μM, MedChemExpress), reversine (1μM, Cayman); hesperadin (0.5μM, Cayman).

### Stable cell lines

Stable isogenic cell lines expressing *hSpindly* constructs were generated by FRT/Flp-In system according to the manufacteŕs intructions (ThermoFisher Scientific). pcDNA5/LAP/hSpindly (a kind gift from Reto Gassman)^12^ or pcDNA5/LAP/hSpindlyT552A were transfected into RPE-1 Flip-In T-Rex cells (a kind gift from Jonathon Pines). After selection in hygromycin, colonies were induced with doxycycline (DOXY) 2μg/mL. pcDNA5/LAP/hSpindlyT552A was constructed using the “Q5 Site Directed Mutagenesis” from New England Biolabs. The sequence of the point mutation was verified on both strands with an automatic sequencer.

### Cell viability assays

Briefly, RPE-1 *GFP-hSpindly* or *RPE-1 GFP-hSpindlyT552A* cells were induced adding 2μg/mL DOXY and arrested in prometaphase with 3.3μM nocodazole for 24 hours. Afterwards, cells were washed out and release of prometaphase in the presence of DOXY. 5×10^3^ or 5×10^2^ cells of each condition were seeded in 6 or 24 well plates, respectively, and incubated for two weeks. Subsequently, colonies were stained with 0.5% crystal violet solution in methanol for 20 minutes, washed, and air-dried. Total stained colonies were decolorated by methanol for 20 minutes. Optic density was quantified at 570nm using a spectrophotometer.

### Immunoblotting

Cells were washed with Phosphate-Buffered Saline (PBS) and lysed in NP40 buffer (150 mM NaCl, 10 mM Tris-HCl (pH 7.5), 1% NP40, 10% glycerol, 1 mM PMSF, 1 μg/mL aprotinin, 1μg/mL pepstatin and 1 μg/mL leupeptin) and incubated on ice for 30 min. After centrifugation at 20,000 *g* for 30 minutes, the supernatant was collected. Concentration was determined using the Bradford assay (Bio-Rad). When necessary, extracts were treated with λ-protein phosphatase (λ-PP)^38^. Proteins were separated by SDS-polyacrilamide gel electrophoresis (SDS-PAGE) and gels were electroblotted onto nitrocellulose membranes and probed with the following antibodies: anti-Bub1 rabbit antibody (1:500, GeneTex GTX30097), anti-BubR1 rabbit antibody (1:1,000, Bethyl Laboratories A300-386A); anti-CyclinB mouse antibody (1:1,000, BD Transduction Lab 610220); anti-β Actin HRP antibody (1:5,000, Santa Cruz sc-47778); anti-GFP rabbit antibody (1:20,000, Immune Systems RGFP-45A); anti-Aurora A kinase rabbit antibody (1:2,000, Novus Biologicals 51843); anti-α tubulin mouse antibody (1:20,000, Sigma T9026); anti-Mad2 rabbit antibody (1:500, Bethyl Laboratories A300-301A); anti-Mad1 mouse antibody (1:500, Santa Cruz sc-47746); anti-hSpindly rabbit antibody (1:1,000, Bethyl Laboratories A301-354A). Anti-phospho-Thr552 hSpindly was generated against the peptide sequence Glu- Arg- Ser- Gly- Asn- pThr- Pro- Asn- Ser- Pro- Arg- Leu corresponding to hSpindly region containing a phosphorylation at threonine 552 (pThr). Peroxidase-coupled donkey anti-rabbit IgG (NA934V) or sheep anti-mouse IgG (NA931) were purchased from GE Healthcare. Immunoreactive bands were visualized using the Enhanced Chemiluminescence (ECL) Western blotting system (GE Healthcare).

### Co-immunoprecipitation experiments

Cell extracts (2mg) were incubated with normal mouse IgG (Santa Cruz, sc-2025) sera for 30 minutes and subsequently with protein G-Sepharose beads (GE Healthcare) for 1 hour at 4 °C. After centrifugation, the beads were discarded, and the supernatants were incubated for 2 hours with anti-Mad1 (Santa Cruz, sc-47746); anti-Mad2 (Santa Cruz, sc-47747) or anti-hSpindly (Novus, H00054908-M01) antibodies, or with normal mouse serum, followed by protein G beads for 1 hour. In case of GFP-Trap agarose beads (ChromoTek), cell extracts were incubated for 2 hours at 4 °C. Beads were washed, and bound proteins were solubilized by adding SDS sample buffer and heated at 95 °C for 5 minutes.

### Fluorescence methods and Microscopy

For fixed-cell imaging, cells were seeded onto glass cover slips (16 mm diameter and thickness number 1.5, 0.16–0.19 mm). Cells were fixed using 3 % paraformaldehyde/4 % sucrose in PBS for 15 minutes. After fixation, cells were washed with PBS and then incubated in permeabilization buffer (0.5 % v/v Triton X-100 in PBS) for 10 minutes. Cells were washed twice with PBS, before 45-60 minutes blocking (3 % Bovine Serum Albumin, 5 % goat serum in 1xPBS). Antibody dilutions were prepared in blocking solution. After blocking, cells were incubated for 1 hour with primary antibody, PBS washed (three washes, 5 minutes each), 1 hour secondary antibody incubation, PBS washed (three washes, 5 min each), mounted with Vectashield containing DAPI (Vector Labs Inc.) and then sealed. The following antibodies were used: mouse anti-Bub1 (1:500, Abcam ab54893); guinea pig anti-CENP-C (1:2,000, PD030 Medical and Biological Labs Company), mouse anti-Mad2 (1:200, Santa Cruz sc-65492), anti-Spindly (1:500, Bethyl Lab A301-354A). For secondary antibodies were used: Alexa Fluor 488, Alexa Fluor 568– or Alexa Fluor 647–conjugated antibody in 3% BSA/PBS (1:500; Invitrogen). Images were captured using in a Leica TCS SP8 microscope with a 1.40 NA 63× objective (HCXPL APO CS2 63/1.40 Oil Leica Microsystems). Using a White Light Laser (WLL). Excitation was performed sequentially at lex 405 nm, 488 nm, 530 nm, 580 nm and 647 nm lasers and emission was captured placing the detectors at Blue, 415/60; Green, 498/540; Orange, 545/85; Red, 595/635; FRed, 662/710. All microscopy data was stored in SACO server in native formats.

### RICS and N&B

Image acquisition was performed in a Leica TCS SP8 microscope with a 1.40 NA 63× objective (HCX PL APO CS2 63/1.40 Oil Leica Microsystems). Using a White Light Laser (WLL), λ_ex_ was set at 488 nm and the band of emission was detected in the range of 498-560 nm via hybrid detectors operating in photon counting mode. 512x512 micrographs were acquired with a pixel size of 50 nm zooming into the sample. Pixel dwell time was set to 10 μs obtaining a line time of 6.24 ms and acquiring one frame in 3.35 s. Analysis was performed using the RICS algorithm of SimFCS program. RICS autocorrelation function has been fitted choosing the model that fits better the data, a diffusion model to calculate the characteristic parameters of the process: the amplitude of the correlation function G0 and the diffusion coefficient D and a binding/unbinding model to calculate the amplitude A (inverse of the number of binding sites) and τ (linearly correlated with the total ‘‘on’’ and ‘‘off’’ time). In order to obtain these parameters, the immobile fraction was subtracted every 4 frames. The excitation volume was calibrated by using a solution of 50 μM of Fluorescein in a solution at pH 9. For this measurement, the scan speed was reduced to 4 μs/pixel. To measure unbinding events in the longer timescale the immobile fraction every 10 frames and 4 frames was calculated, and their difference was analyzed. On the same measurements for RICS, N&B analysis was performed. In order to calibrate the brightness, cells expressing monomeric GFP were analyzed to determine the brightness of the monomer.

## ETHICS DECLARATIONS

### Competing interests

The authors declare no competing interests.

## AUTHOR CONTRIBUTIONS

Conceptualization: M.M.-S; Methodology: M.M.-S., J.H.-R., D.F.-G., A.B.-F., N.F., F.R.; Software: N.F.; Validation: M.M.-S, N.F., F.R.; Formal analysis: M.M.-S, D.F.-G, N.F.; Investigation: M.M.-S, J.H.-R, D.F.-G., A.B.-F., N.F.; Resources: M.M.-S, N.F., F.R.; Writing-original draft: M.M.-S; Writing-review & editing: M.M.-S., J.H.-R., D.F.-G., A.B.-F., N.F., F.R.; Visualization: M.M.-S, D.F.-G, N.F.; Supervision: M.M.-S; Funding acquisition: M.M.-S, N.F., F.R.

## ACKNOWLEDGEMENTS

M.M.-S. was a Marie-Curie fellow funding by Excellence Science Marie Sklodowska-Curie Actions grant agreement 101024268.

This work was supported by grants PID2023-152247NB-C21 and PID2022-140551NB-100 funded by Ministerio de Ciencia, Innovación y Universidades.

We thank to Nina Pucekova (Centre of Biomedical Sciences, Warwick Medical School, University of Warwick, Coventry, UK) for critical reading of the manuscript.

## Notes

### Competing Interest Statement

The authors have declared no competing interest.

## REFERENCES

1. McAinsh, A. D. & Kops, G. J. P. L. Principles and dynamics of spindle assembly checkpoint signalling. Nat Rev Mol Cell Biol 24, 543–559 (2023).

2. Musacchio, A. The Molecular Biology of Spindle Assembly Checkpoint Signaling Dynamics. Current Biology 25, 1002–1018 (2015).

3. Cmentowski, V. et al. RZZ-Spindly and CENP-E form an integrated platform to recruit dynein to the kinetochore corona. EMBO J 42, e114838 (2023).

4. Raisch, T. et al. Structure of the RZZ complex and molecular basis of Spindly-driven corona assembly at human kinetochores. EMBO J 41, e110411 (2022).

5. Silió, V., McAinsh, A. D. & Millar, J. B. KNL1-Bubs and RZZ Provide Two Separable Pathways for Checkpoint Activation at Human Kinetochores. Dev Cell 35, 600–613 (2015).

6. Ying, W. C. et al. Mitotic control of kinetochore-associated dynein and spindle orientation by human Spindly. Journal of Cell Biology 185, 859–874 (2009).

7. Griffis, E. R., Stuurman, N. & Vale, R. D. Spindly, a novel protein essential for silencing the spindle assembly checkpoint, recruits dynein to the kinetochore. Journal of Cell Biology 177, 1005–1015 (2007).

8. Barisic, M. et al. Spindly/CCDC99 is required for efficient chromosome congression and mitotic checkpoint regulation. Mol Biol Cell 21, 1968–1981 (2010).

9. Maciejowski, J. et al. Mps1 Regulates Kinetochore-Microtubule Attachment Stability via the Ska Complex to Ensure Error-Free Chromosome Segregation. Dev Cell 41, 143–156 (2017).

10. Moudgil, D. K. et al. A novel role of farnesylation in targeting a mitotic checkpoint protein, human spindly, to kinetochores. Journal of Cell Biology 208, 881–896 (2015).

11. Sacristan, C. et al. Dynamic kinetochore size regulation promotes microtubule capture and chromosome biorientation in mitosis. Nat Cell Biol 20, 800–810 (2018).

12. Gassmann, R. et al. Removal of Spindly from microtubule-attached kinetochores controls spindle checkpoint silencing in human cells. Genes Dev 24, 957–971 (2010).

13. Silva, P. M. A. et al. Suppression of spindly delays mitotic exit and exacerbates cell death response of cancer cells treated with low doses of paclitaxel. Cancer Lett 394, 33–42 (2017).

14. Tian, X. & Wang, N. Upregulation of ASPM, BUB1B and SPDL1 in tumor tissues predicts poor survival in patients with pancreatic ductal adenocarcinoma. Oncol Lett 19, 3307–3315 (2020).

15. Yamano, H. APC/C: current understanding and future perspectives. F1000Research 8, 10.12688/f1000research.18582.1 (2019).

16. Alfieri, C., Zhang, S. & Barford, D. Visualizing the complex functions and mechanisms of the anaphase promoting complex/cyclosome (APC/C). Open Biology 7, 10.1098/rsob.170204 (2017).

17. Currie, C. E., Mora-Santos, M., Smith, C. A., McAinsh, A. D. & Millar, J. B. A. Bub1 is not essential for the checkpoint response to unattached kinetochores in diploid human cells. Current Biology vol. 28 R929–R930 (2018).

18. O’Connor, A. et al. Requirement for PLK1 kinase activity in the maintenance of a robust spindle assembly checkpoint. Biol Open 5, 11–19 (2016).

19. Huang, H. et al. Phosphorylation sites in BubR1 that regulate kinetochore attachment, tension, and mitotic exit. Journal of Cell Biology 183, 667–680 (2008).

20. Rodriguez-Rodriguez, J. A. et al. Distinct Roles of RZZ and Bub1-KNL1 in Mitotic Checkpoint Signaling and Kinetochore Expansion. Current Biology 28, 3422–3429 (2018).

21. Zhang, G. et al. Efficient mitotic checkpoint signaling depends on integrated activities of Bub1 and the RZZ complex . EMBO J 38, e100977 (2019).

22. Baron, A. P. et al. Probing the catalytic functions of Bub1 kinase using the small molecule inhibitors BAY-320 and BAY-524. Elife 5, e12187 (2016).

23. Garcia, Y. A. et al. Mapping Proximity Associations of Core Spindle Assembly Checkpoint Proteins. J Proteome Res 20, 3414–3427 (2021).

24. Siemeister, G. et al. Inhibition of BUB1 kinase by Bay 1816032 sensitizes tumor cells toward taxanes, ATR, and PARP inhibitors in vitro and in vivo. Clinical Cancer Research 25, 1404–1414 (2019).

25. Van Vuuren, R. J., Visagie, M. H., Theron, A. E. & Joubert, A. M. Antimitotic drugs in the treatment of cancer. Cancer Chemotherapy and Pharmacology 76, 1101– 1112, 10.1007/s00280-015-2903-8 (2015).

26. Bolanos-Garcia, V. Assessment of the Mitotic Spindle Assembly Checkpoint (SAC) as the Target of Anticancer Therapies. Curr Cancer Drug Targets 9, 131–141 (2009).

27. Bargiela-Iparraguirre, J. et al. Mad2 and BubR1 modulates tumourigenesis and paclitaxel response in MKN45 gastric cancer cells. Cell Cycle 13, 3590–3601 (2014).

28. Gurden, M. D. et al. Naturally occurring mutations in the MPS1 gene predispose cells to kinase inhibitor drug resistance. Cancer Res 75, 3340–3354 (2015).

29. Kurai, M. et al. Expression of Aurora kinases A and B in normal, hyperplastic, and malignant human endometrium: Aurora B as a predictor for poor prognosis in endometrial carcinoma. Hum Pathol 36, 1281–1288 (2005).

30. Digman, M. A. et al. Measuring fast dynamics in solutions and cells with a laser scanning microscope. Biophys J 89, 1317–1327 (2005).

31. Yamamoto, T. G., Watanabe, S., Essex, A. & Kitagawa, R. Spdl-1 functions as a kinetochore receptor for MDF-1 in Caenorhabditis elegans. Journal of Cell Biology 183, 187–194 (2008).

32. Allan, L. A. et al. Cyclin B1 scaffolds MAD 1 at the kinetochore corona to activate the mitotic checkpoint . EMBO J 39, e103180 (2020).

33. Maldonado, M. & Kapoor, T. M. Constitutive Mad1 targeting to kinetochores uncouples checkpoint signalling from chromosome biorientation. Nat Cell Biol 13, 475–482 (2011).

34. Gama, J. B. et al. Molecular mechanism of dynein recruitment to kinetochores by the Rod-Zw10-Zwilch complex and Spindly. Journal of Cell Biology 216, 943–960 (2017).

35. McKenney, R. J., Huynh, W., Tanenbaum, M. E., Bhabha, G. & Vale, R. D. Activation of cytoplasmic dynein motility by dynactin-cargo adapter complexes. Science (1979) 345, 337–341 (2014).

36. D’amico, E. A. et al. Conformational transitions of the Spindly adaptor underlie its interaction with Dynein and Dynactin. Journal of Cell Biology 221, e202206131 (2022).

37. Belmonte-Fernández, A. et al. Cisplatin-induced cell death increases the degradation of the MRE11-RAD50-NBS1 complex through the autophagy/lysosomal pathway. Cell Death Differ 30, 488–499 (2023).

38. Mora-Santos, M. et al. Glycogen synthase kinase-3β (GSK3β) negatively regulates PTTG1/human securin protein stability, and GSK3β inactivation correlates with securin accumulation in breast tumors. Journal of Biological Chemistry 286, 30047–30056 (2011).

