## Supplementary Information for "hSpindly’s dynamic controls SAC activity independently of the KBB pathway at unattached kinetochores"

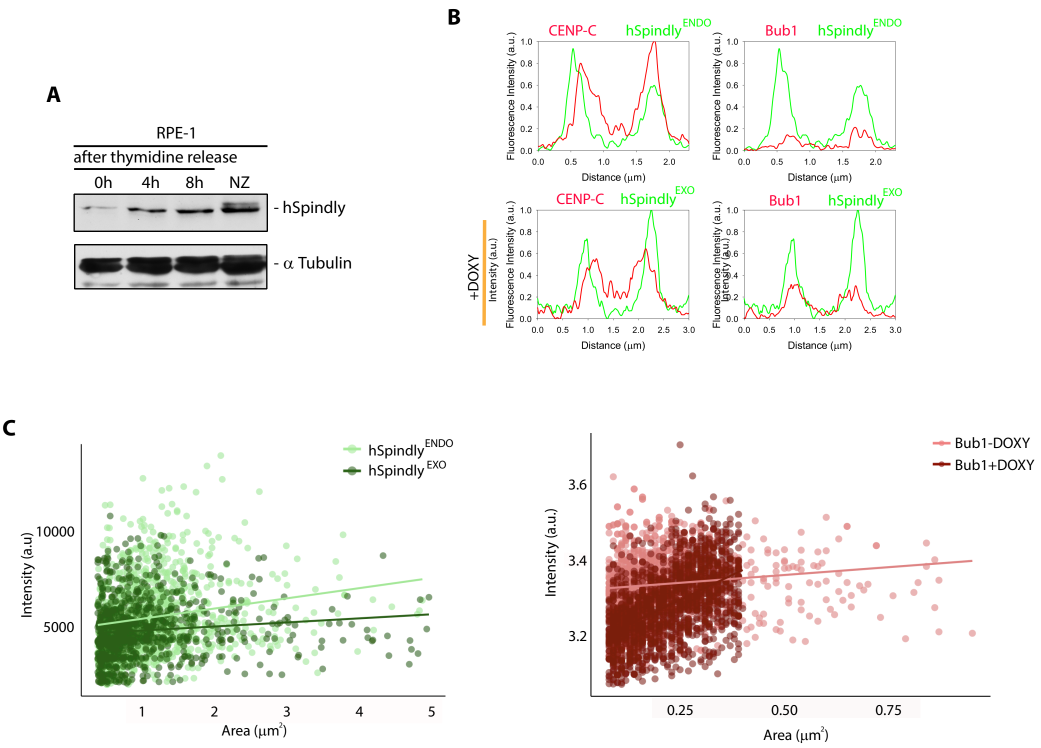


**Supplemental 1, related to Figure 1**

**A.** RPE-1 cells were synchronized at the G1/S boundary using 2.5 mM thymidine and either collected at indicated time points or treated with NZ for 24 hours. Cell extracts were immunoblotted with the indicated antibodies. **B.** Intensity profile of $hSpindly^{ENDO}$ or $hSpindly^{EXO}$ and CENP-C or Bub1, in the absence or presence of DOXY in the cross-section of a kinetochore from mitotic cells. **C.** Distribution of $hSpindly^{ENDO}$ or $hSpindly^{EXO}$ populations based on kinetochore signal intensity and area. Each dot represents a kinetochore; lines indicate the mean.

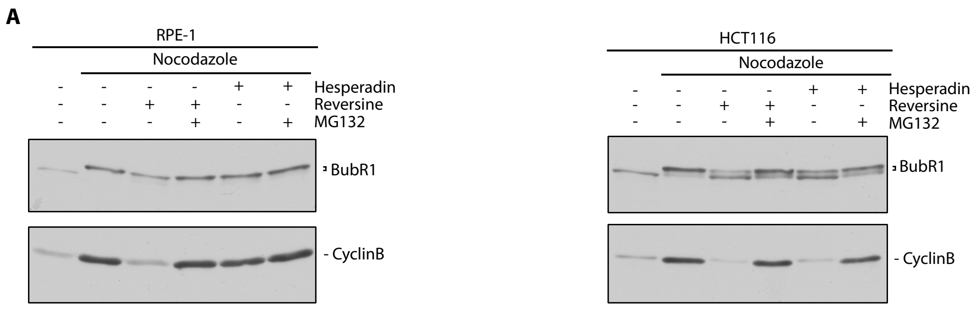


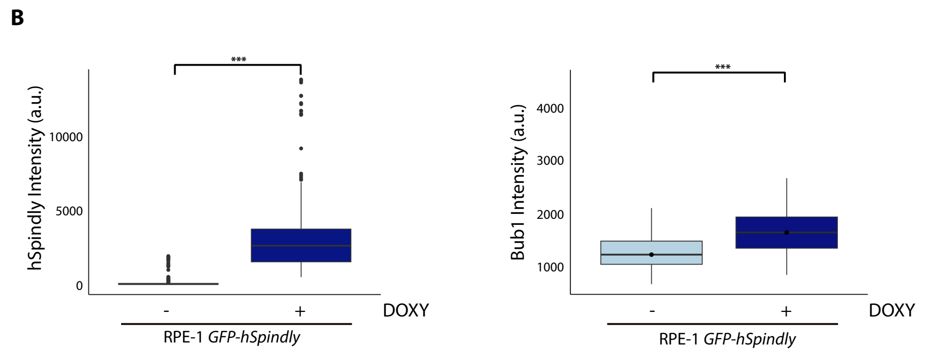


**Supplemental Figure 2, related to Figure 2**

**A.** RPE-1 (panel left) or HCT116 (panel right) were treated with 3.3µM NZ for 24 hours. Before collecting, mitotic cells were incubated with 0.5µM hesperadin or 1µM reversine in the presence of proteasome inhibitor (MG132) 20µM for 90 minutes. Western-blots of extracts were analyzed for BubR1 and Cyclin B levels. **B.** Box plots representing levels of GFP-hSpindly or Bub1 at kinetochores in the presence or absence of DOXY. Statistical analysis was performed with a nonparametric t-test comparing two unpaired groups (***, p<0.001).

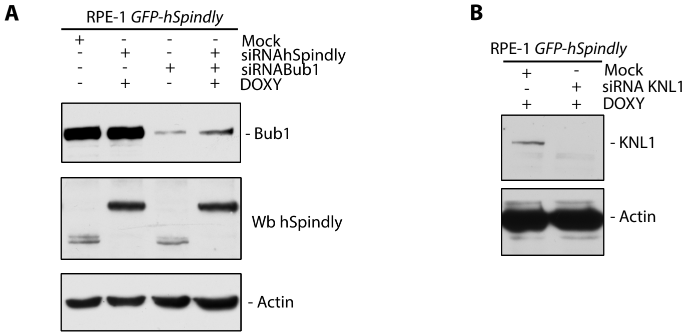


**Supplemental Figure 3, related to Figure 3**

**A.** RPE-1 *GFP-hSpindly* cells were transfected with siRNAhSpindly or siRNABub1 for 48 hours, and treated with NZ in the presence or absence of DOXY for 24 hours before harvesting. Extracts were immunoblotted with the indicated antibodies. In Wb hSpindly, GFP-hSpindly is the upper band and hSpindly endogenous is the lower band. **B.** RPE-1 *GFP-hSpindly* prometaphase cells were treated with the indicated siRNAs, and extracts were blotted with the indicated antibodies.

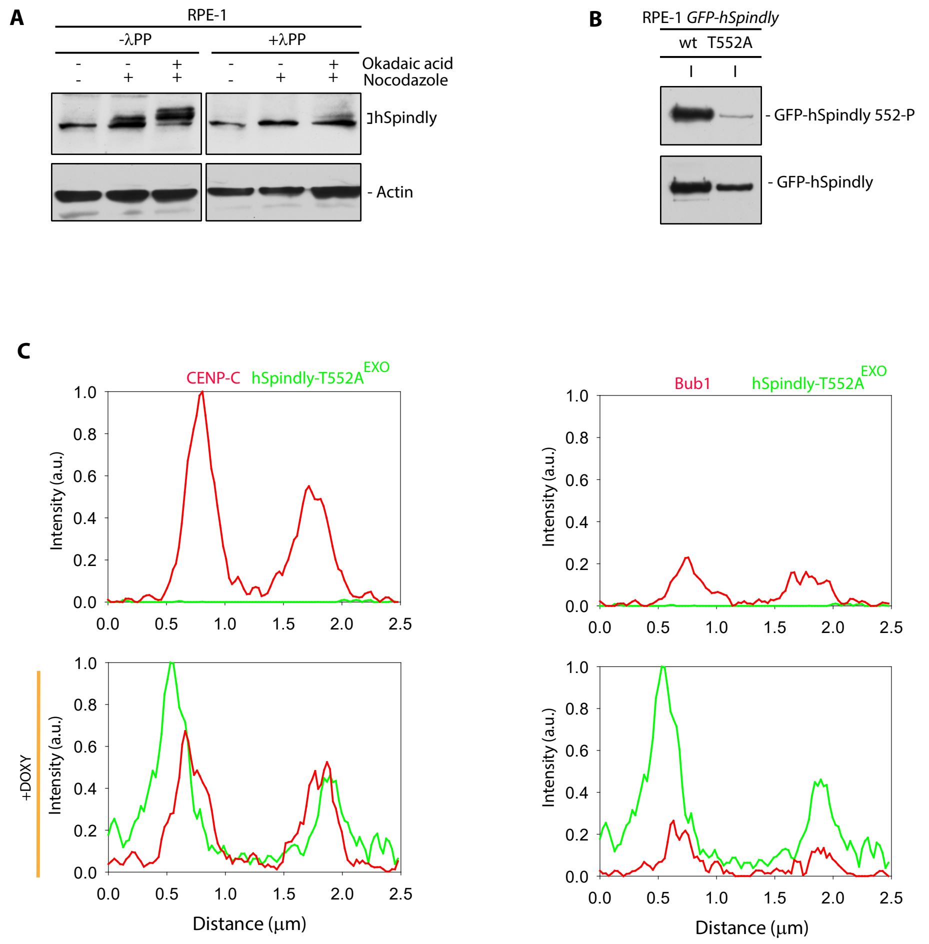


**Supplemental Figure 4, related to Figure 4**

**A.** RPE-1 cells were treated or not with NZ for 24 hours or 1µM okadaic acid for 90 minutes. Indicated extracts were treated with λPP for 30 minutes at 37ºC and immunoblotted with indicated antibodies. **B.** RPE-1 *GFP-hSpindly* or *RPE-1 GFP-hSpindly T552A* prometaphase cell extracts induced previously by DOXY addition were immunoprecipitated using GFP-Trap beads. Anti-phospho-Thr552 hSpindly was used for immunoblotting. I: Immune. **C.** Intensity profile of hSpindly T552A and CENP-C or Bub1, in the absence or presence of DOXY in the cross-section of a kinetochore from mitotic cells.

**A**

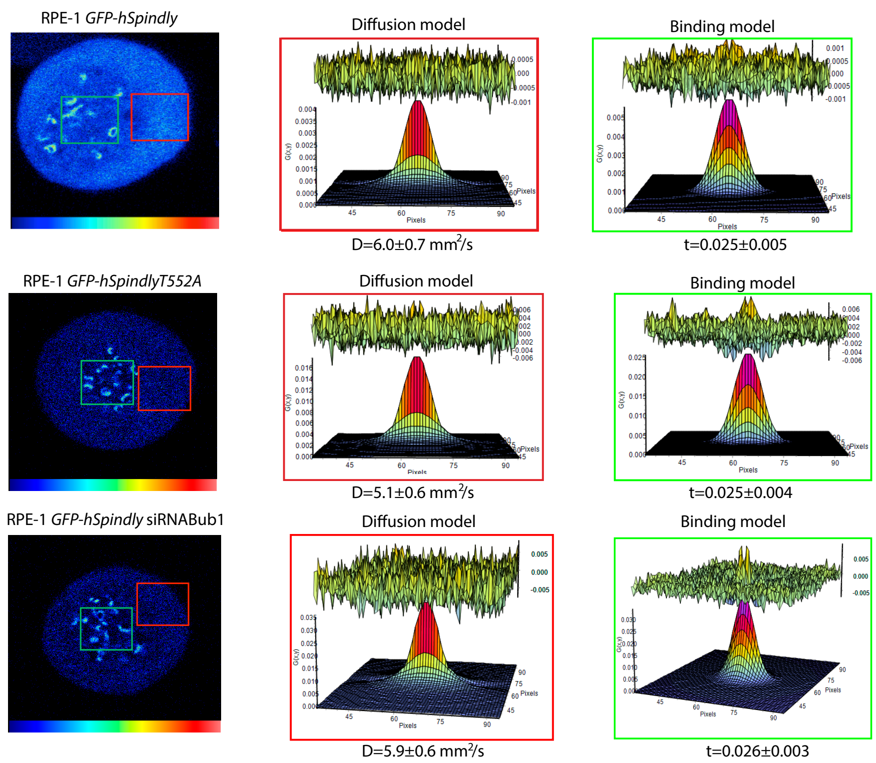


**Supplemental Figure 5, related to Figure 6**

**A.** 512x512 fluorescence intensity micrographs (left column) of RPE-1 cells transfected with GFP-hSpindly (top row), GFP-hSpindly-T552 (middle row) or GPF-hSpindly in siRNA Bub1 background (bottom row). The red and green ROIs indicate where the spatial correlation functions were fitted for the diffusion (middle column) and binding model (right column), respectively.

**Supplemental Movies related to Figure 6**

**Supplemental Movie 1.** RPE-1 *GFP-hSpindly* cells treated with NZ and DOXY for 24 hours.

**Supplemental Movie 2.** RPE-1 *GFP-hSpindlyT552A* cells treated with NZ and DOXY for 24 hours.

**Supplemental Movie 3.** RPE-1 *GFP-hSpindly* cells treated with siRNA Bub1 for 48 hours and treated with NZ and DOXY for 24 hours before recording.
